# The transcription factor Ste12(1) from *Nakaseomyces glabratus* acts pheromone-responsive in *S. cerevisiae* but seems decoupled from its cognate upstream GPCR in its native host

**DOI:** 10.64898/2026.09.17.752268

**Authors:** Min Lu, Thijs de Vroet, Sonja Billerbeck

## Abstract

G-protein-coupled receptor (GPCR)-mediated signalling pathways govern critical cellular responses in fungi, including mating, biofilm formation, and virulence. The mating pathway of *Saccharomyces cerevisiae*, in which peptide-sensing GPCRs (peptide GPCRs) activate a MAP kinase cascade and the transcription factor Ste12, has thereby served as the canonical model for fungal peptide GPCR signalling. Homologues of this pathway are conserved across many fungi, including the high-priority pathogen *Nakaseomyces glabratus*. Yet the function of this pathway in this mating-incompetent pathogen remains unclear. Here, we investigated whether the central transcription factor Ste12 of *N. glabratus* (NgSte12(1)) can act as a peptide GPCR-coupled transcription factor using a component-swapping strategy in which pathway elements from *N. glabratus* and *S. cerevisiae* were reciprocally exchanged and assessed using fluorescent and growth arrest readouts. We show that NgSte12(1) is in principle capable of supporting peptide-GPCR-dependent activation in *S. cerevisiae*, however, subtle and dependent on promoter identity. Ste12 chimaeras further demonstrate that all three NgSte12(1) subdomains are individually functional and compatible with their *S. cerevisiae* counterparts. Heterologous expression of ScSte12 in *N. glabratus* shows constitutive pathway activation, while NgSte12(1) does not show pathway activation in *N. glabratus*. Peptide-GPCR expression experiments indicate a block in *N. glabratus* signalling upstream of Ste12 (1): the *S. cerevisiae* peptide-GPCR Ste2 (ScSte2) activated the *N. glabratus* Ste12 GPCR-dependent downstream signalling constitutively, while NgSte2 failed to activate signalling in its native environment. Together, these results indicate that NgSte12(1) is a competent pheromone-responsive transcription factor that can couple to a heterologous GPCR in a heterologous context but is unresponsive to its own cognate upstream peptide/GPCR pair, either because it binds a yet unknown ligand or the signalling is rewired in this pathogen.

## INTRODUCTION

G-protein-coupled receptors (GPCRs) are a large and diverse family of seven-transmembrane domain receptors that transduce extracellular signals into intracellular responses. Together with their downstream signalling components, GPCRs represent one of the most widespread sensory systems used by eukaryotes to perceive and respond to their environment [1]. In humans, GPCR-mediated signalling is implicated in virtually all major disease areas, and approximately 35% of all approved drugs exert their therapeutic effects by targeting these receptors [2].

Fungi likewise encode a broad repertoire of GPCRs [3]; however, for the majority of fungal GPCRs, the cognate ligands, downstream signalling cascades, and associated cellular responses remain poorly characterised. Accumulating evidence nonetheless points to critical roles for fungal GPCR signalling in virulence, both in plant and human pathogenic species [3, 4]. Given the urgent global need for novel antifungal strategies to combat the growing burden of drug-resistant fungal pathogens [5], a deeper understanding of these signalling systems is of considerable importance, as they have been suggested as future drug targets [3].

One fungal GPCR system that is exceptionally well characterised is the canonical mating pathway of *Saccharomyces cerevisiae*, which has served as a valuable model framework for basic research and engineering for decades [6–11]. This pathway relies on two peptide pheromone-sensing GPCRs (hereafter short peptide-GPCRs), Ste2 and Ste3, coupled to a mitogen-activated protein kinase (MAPK) cascade, eventually leading to the activation of the transcription factor Ste12. Ste12 eventually regulates the expression of over 200 mating-associated genes [12]. Importantly, in several human and plant fungal pathogens, homologues of this mating pathway have been shown to govern behaviours beyond mating itself, including biofilm formation and virulence [13–15]. In this context, several fundamental questions about signal perception and transduction remain unanswered: for example, whether the two mating GPCRs Ste2 and Ste3 are capable of sensing alternative ligands beyond their cognate peptide pheromones, whether the cell can discriminate between mating and non-mating signals and responses, and if so, by what molecular mechanisms.

The human fungal pathogen *Nakaseomyces glabratus* presents a particularly compelling case in this context. This organism encodes clear homologues of both mating GPCRs (Ste2 and Ste3), and the full MAPK signalling cascade [16] **(Supplementary Table 1)**. Differences are that it encodes two copies of the transcription factor Ste12 (called Ste12(1) and (2) [17]), while a homologue for Dig2, a negative regulator of Ste12 seems missing. Notably, a Dig2 homologue is also missing in *C. albicans* [18]. Functional conservation of the pathway’s core components is further supported by previous findings that both NgSte12 variants, as well as the protein kinases NgSte11, and NgSte20 can each complement their corresponding mating-defective deletions in *S. cerevisiae* [17, 19–21]. Interestingly, mating has never been directly observed in *N. glabratus* under laboratory conditions [16, 22], while genomic evidence is consistent with occasional sexual reproduction [16, 23]. Mating events may thus be rare or restricted to highly specialised environments, as has been demonstrated for *S. cerevisiae* in the wasp gut [24]. This raises the fundamental question of whether the mating pathway is functionally active in *N. glabratus* and, if so, whether it serves a biological role beyond mating. Therefore, throughout this article, we refer to the ‘mating pathway’ as peptide-GPCR downstream signalling in *N. glabratus*, as its role in mating remains unclear.

Addressing this question carries significant clinical relevance. *N. glabratus* is an opportunistic human pathogen and the second most common causative agent of candidemia worldwide, accounting for approximately 5-30% of *Candida* isolates from infected patients, with a steadily increasing incidence [25]. The species exhibits rising resistance to multiple antifungal drug classes and has been designated a high-priority fungal pathogen by the World Health Organization (WHO) [26]. Formerly classified as *Candida glabrata*, it has since been reclassified into the genus *Nakaseomyces* [27]. Several components of the mating-like MAPK pathway in *N. glabratus*, including Ste12, Ste11, Ste20, and Fus3, have already been functionally linked to pathogenesis [28], suggesting that this signalling pathway may be co-opted for virulence-related functions. Understanding how GPCR signalling and downstream pathway activation operate in this pathogen could therefore reveal novel targets for antifungal intervention.

Here, we begin to dissect this pathway by focusing on the central transcription factor Ste12. We investigated whether it can be activated by an upstream peptide-GPCR upon peptide ligand binding. While it was shown before that plasmid-based expression of both of *N. glabratus’* NgSte12s could functionally complement an Ste12 deletion in *S. cerevisiae* [19], whether this was constitutive or dependent on peptide-GPCR activation had not been examined. It was also unclear if the NgSet12 transcription factor could be activated by a peptide-GPCR in its native host *N. glabratus*.

We focussed on one of the two NgSte12s, specifically NgSte12(1) as a representative candidate: Ste12(1) and Ste12(2) share 47% sequence conservation between each other. Furthermore, both NgSte12s contain a conserved canonical Ste12 DNA-binding motif, with an F-to-Y substitution in Ste12(2), and both lack the conserved Dig1-binding motif in their pheromone response domain **(Supplementary Figure 1 and 2)**.

While the peptide-GPCR(Ste2)/Ste12 activation cascade is well characterised in *S. cerevisiae* [29–31], functionality in *N. glabrata* remains uncertain. Based on this, we employed a component-swapping approach **(Figure 1A)** to investigate whether a functional peptide-GPCR/Ste12/reporter cascade could be reconstituted through the reciprocal exchange of pathway components between the two species, specifically the peptide-GPCR and the transcription factor Ste12. We extended this swapping strategy to individual Ste12 subdomains, encompassing the DNA-binding domain (DBD), pheromone response domain (PRD), and transcriptional activation domains (AD), and tested the resulting chimeras in *S. cerevisiae*.

**Figure 1:**
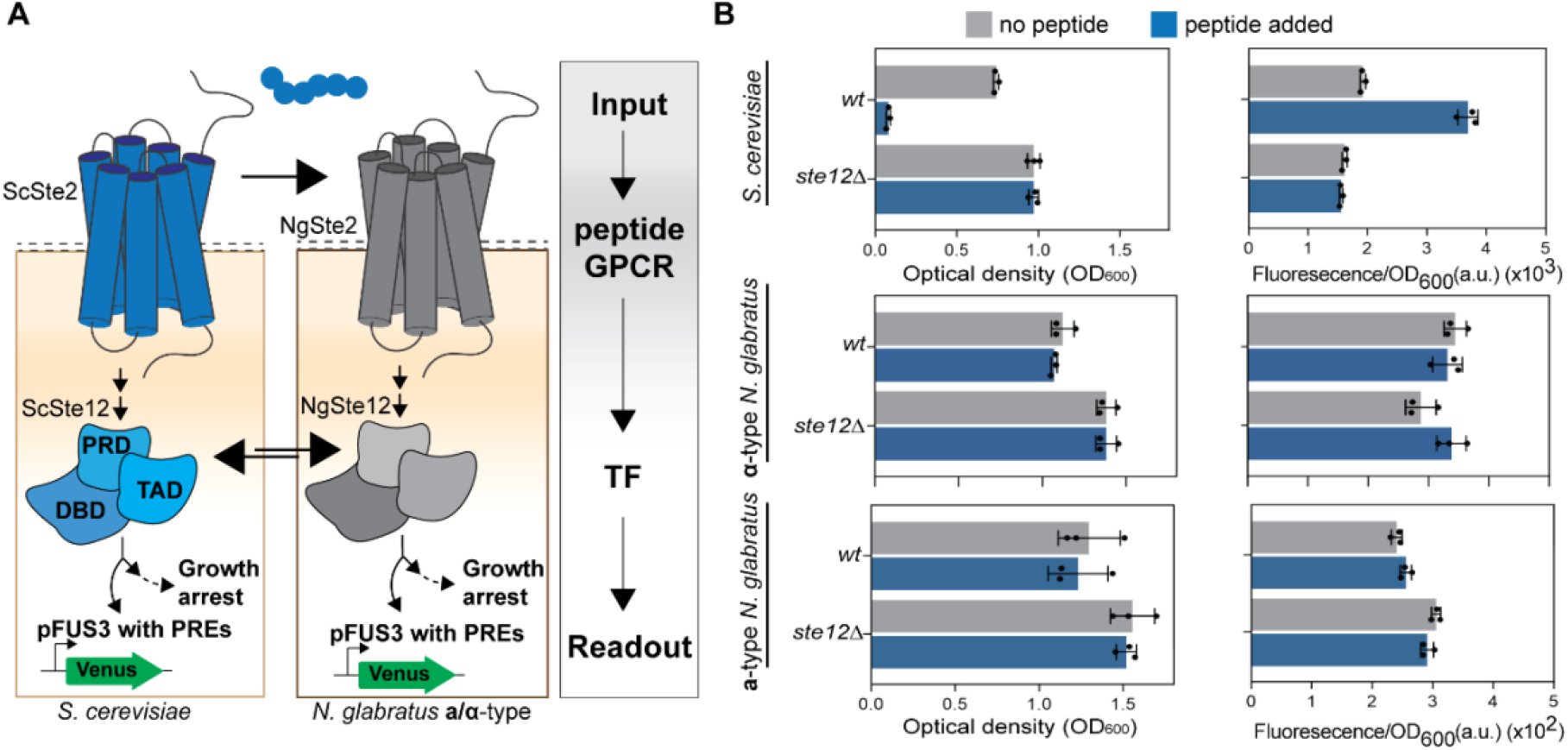
Peptide-GPCR-Ste12 signalling pathway in both *S. cerevisiae* and *N. glabratus*. **A**. Schematic overview on the component swapping approach to study functionality of the peptide-GPCR/Ste12/readout cascade in *N. glabratus*. NgSte12(1) and its three subdomains were heterogeneously expressed in *S. cerevisiae* and tested for functionality. ScSte12 and the peptide GPCR Ste2 were heterogeneously expressed in both mating types of *N. glabratus*. Growth arrest (OD_600_) and activation of a pheromone response element (PRE)-containing promoter linked to a fluorescent reporter was used as a readout. Note: Functional heterologous expression of the *N. glabratus* peptide-GPCR NgSte2 in *S. cerevisiae* was studied before and therefore not included here [32]. **B**. Phenotypes of the six test strains in the presence or absence of 10 µM peptide ligand. Both readouts were assayed by measuring OD_600_ (cell cycle arrest) and fluorescent activation (activation of PRE-containing pFUS3 promoter) after 20 hours of incubation at 30°C. Data represent means ± SD from three biological replicates. DBD: DNA binding domain. PRD: Pheromone response domain. AD: activation domain. PRE: pheromone response element.

The rationale underlying this component-swapping approach was that the establishment or disruption of a chimeric cascade assembled from mixed-species components could help pinpoint which element failed to function in a heterologous context.

We used a fluorescent reporter and growth inhibition via cell cycle arrest as functional readouts **(Figure 1A)**, and *S. cerevisiae* (MATa, with and without *ste12* deleted) and *N. glabratus* (MATa and MATα, each with and without *ste12(1)* deleted) as host strains.

## RESULTS AND DISCUSSION

### Experimental design

To enable the swapping, six core strains were used **(Supplementary Table 2)**: The available *S. cerevisiae ste12* deletion strain BY4733 *ste12Δ::meth1* (MATa) **(strain 1)** [33] served as a well-defined heterologous background for Ste12 characterisation in a MATa background. This strain is capable of alpha-factor pheromone (called peptide hereafter) sensing via its endogenous peptide-GPCR Ste2. BY4741 without *ste12* deleted served as a control **(strain 2)**. Two *N. glabratus* strains of opposite mating types were used to characterise NgSte12 and ScSte12 in *N. glabratus* and to evaluate potential mating-type-dependent effects: ATCC2001 HTL^−^ *ste12(1)Δ::nat* (MATα) **(strain 3)** and ySB38 *his3*Δ::*frt trp1*Δ::*frt ste12(1)*Δ::*nat* (MATa) **(strain 4)**. The mating types of both *N. glabratus* strains were verified by PCR **(Supplementary Figure 3)**. The MATα strain should, in theory, not be able to sense its cognate alpha-pheromone peptide, as transcriptomics data show it expresses the a-factor-sensing GPCR Ste3 (**Supplementary Table 3** [33]). However, another study reported comparable transcription levels of Ste2 and Ste3, raising the possibility that Ste3 may also contribute to α-pheromone sensing [16]. ATCC2001 HTL^−^ and ySB38 *his3*Δ::*frt trp1*Δ::*frt* without *ste12* deletions served as controls **(strains 5 and 6**).

The *N. glabratus* MATα *ste12* deletion strain was generated from the well-characterised ATCC2001 HTL^−^ strain [34] by homologous recombination and verified by full genome sequencing **(Supplementary Figure 4** and **Supplementary Table 4)**. The MATa strain ySB38 first required additional engineering to make it genetically tractable: the auxotrophic markers *his3* and *trp1* were deleted to enable selection of exogenously transformed plasmids. This was done by homologous recombination using a Nourseothricin (Nat) marker for selection and using a FLP/FRT system to allow recycling of the Nat selectable marker [35], followed by *ste12(1)* deletion via CRISPR/Cas9. All genomic modifications were verified by Illumina-based full genome sequencing **(Supplementary Figure 4 and Supplementary Table 4)**.

For Ste12 swapping, ScSte12 and NgSte12(1) were each expressed under the control of their respective native promoters, from the low-copy vector pRS413, generating plasmids P_ScSte12_-ScSte12 and P_NgSte12_-NgSte12. The pRS413 vector without insert served as a control (P _Empty Vector1_) **(Supplementary Table 5 and 6)**.

A fluorescence readout using the pheromone-responsive NgFUS3 promoter linked to a Venus reporter in the low-copy vector pRS414 was used to test for transcriptional promoter activation via the pheromone response element (PRE, or Ste12 binding site), a widely used proxy for GPCR activation in yeast [36]. In parallel, growth inhibition (a proxy for Far1-induced cell-cycle arrest) was measured using optical density at 600 nm (OD_600_) as a secondary functional readout, related to the more complex mating response. A pRS414 vector without inserts served as a control (P _Empty Vector2_) for the Venus reporter to study NgSte12(1) and ScSte12 promoter strength.

The peptide-GPCRs ScSte2 and NgSte2 were each cloned under the control of the TDH3 promoter and ENO1 terminator into the low-copy vector pRS413, generating plasmids P_TDH3-_ScSte2 and P_TDH3_-NgSte2. The promoter-terminator combination had previously been shown to support functional Ste2 complementation [32]. GPCR activation was assessed using 10 µM *S. cerevisiae* mating peptide (sequence: WHWLQLKPGQPMY), as this concentration had previously been shown to activate both, ScSte2 and NgSte2 [32]. The latter to a comparable extent to the *N. glabratus*-endogenous mating peptide (sequence: WHWVRLRKGQGLF) [32].

### Full-length NgSte12(1) and all its sub-domains support peptide-GPCR-dependent activation in *S. cerevisiae*, however full-length NgSte12(1) only subtle and in dependence on promoter context

We first tested if the six test strains carrying the Venus reporter plasmid and the empty vector pRS413 behaved as expected. Wild-type BY4741 showed growth arrest and fluorescence activation upon peptide treatment **(Figure 1B)**, while both responses were abolished in BY4733 *ste12Δ*; the four *N. glabratus* wild-type and *Ste12* deletion strains showed no peptide-responsiveness. **(Figure 1B)**. Notably, basal fluorescence from the FUS3 promoter was about 5-fold lower in both *N. glabratus* mating types when compared to *S. cerevisiae* **(Figure 1B)**.

We then started with characterising NgSte12(1) in *S. cerevisiae* by transforming BY4733 *ste12Δ* with P_NgSte12_-NgSte12(1) and the Venus reporter and assessing growth arrest and fluorescence activation in the presence and absence of 10 µM peptide: interestingly, NgSte12(1) induced growth arrest even in the absence of pheromone **(Figure 2A)**, indicating constitutive activation of the mating pathway. Constitutive growth arrest thereby manifested as slow colony growth after transformation, slow growth in culture, and strain instability (meaning the phenotype was lost upon prolonged maintenance on agar plates). To confirm that growth arrest was Ste12(1)-dependent rather than a result of metabolic burden, we deleted *far1*, a key mediator of pheromone-induced G1 arrest. BY4733 *ste12Δfar1Δ* cells expressing NgSte12(1) could grow normally **(Supplementary Figure 5)**, confirming the involvement of Far1 in the NgSte12(1)-induced phenotype. In the fluorescence assay, however, NgSte12(1) only marginally elevated Venus reporter expression compared with the *ste12Δ* strain **(Figure 2A)**, with no further increase upon peptide treatment. This suggested that while NgSte12(1) drove constitutive Far1-dependent cell cycle arrest, it did not activate pheromone-responsive transcription from the FUS3 promoter. It could also mean that cells were too sickened by the constitutive growth arrest phenotype to respond.

**Figure 2:**
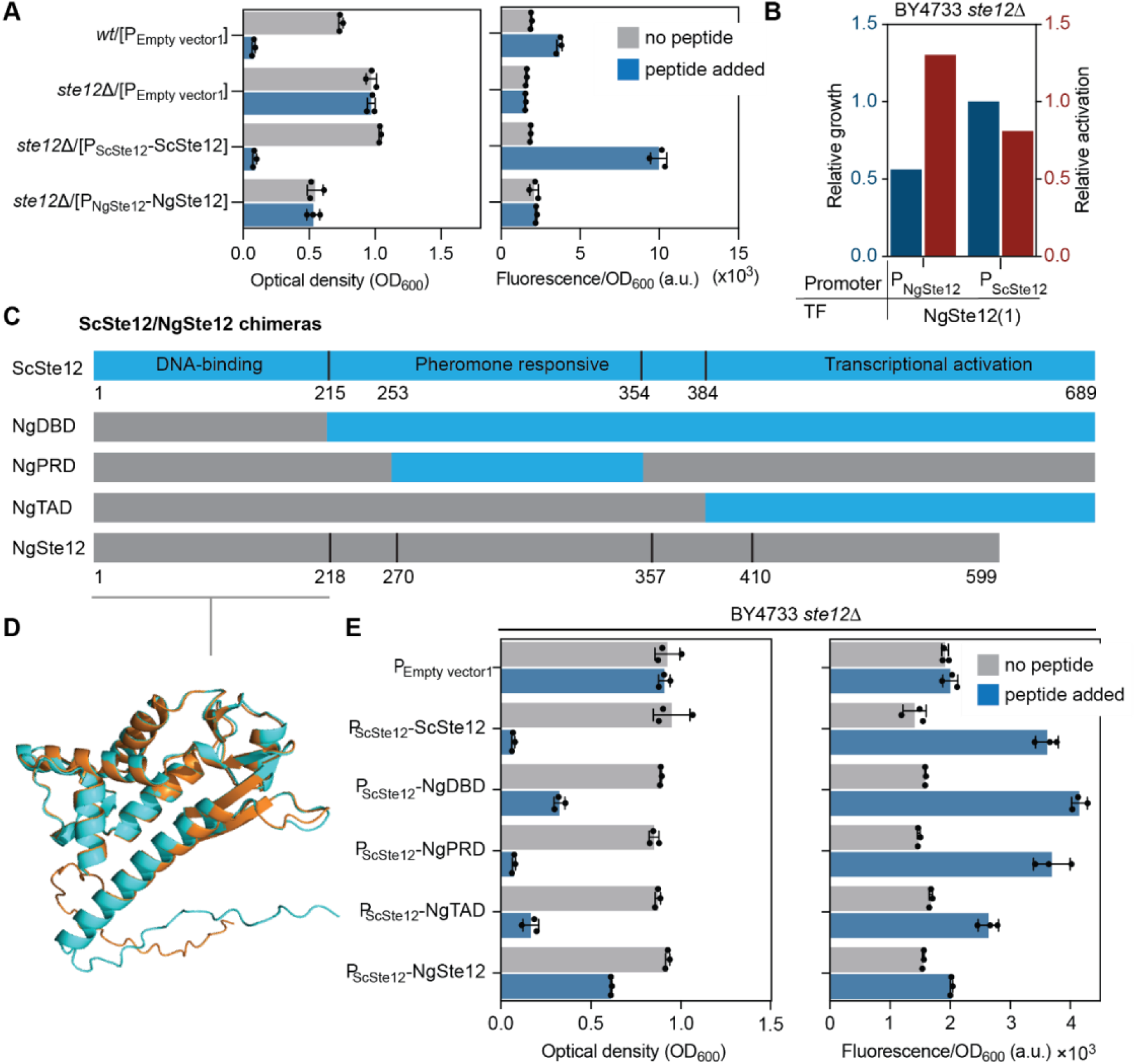
Expression of full-length NgSte12 and Sc/NgSte12 chimeras in *S. cerevisiae*. **A**. Ste12-expressing plasmids empty vectors were used to transform *S. cerevisiae* wild-type or *ste12*Δ strains. ScSte12 and NgSte12(1) were expressed from their respective native promoters. All strains were co-transformed with a Venus reporter plasmid. Growth was assessed by optical density at 600 nm (OD_600_). Ste12-dependent transcriptional activity was monitored using fluorescence normalized to OD_600_. Strains were tested in the presence or absence of 10 µM peptide. Data represent means ± SD from three biological replicates. **B**. Comparative graphing of Ste12-induced growth arrest and fluorescent activation in the absence of peptide (thus constitutive) when NgSte12(1) was expressed from its own promoter (P_NgSte12_) or the *S. cerevisiae*-derived Ste12 promoter (P_ScSte12_). Data are shown in relation to the phenotype of wild-type cells, which was set to 1 (100%). **C**. Ste12 domain organization and outline of Sc/NgSte12 chimeras. ScSte12 comprises three functional domains: the DNA-binding domain (DBD), the pheromone-responsive domain (PRD), and the transcriptional activation domain (TAD). ScSte12 domains were replaced with corresponding regions from NgSte12(1). Alignments used to extract domain boundaries are shown in **Supplementary Figure 1**. The amino acid sequences of the chimeras are listed in **Supplementary Table 7. D**. Conservation comparison of DBD structures from ScSte12 (1-215aa, orange) and NgSte12 (1-218aa, light blue). Their structures were predicted with AlphaFold 3 [42] and aligned using PyMol [43–45]. **E**. Ste12 chimeras and full-length Ng and ScSte12 were expressed from the ScSte12 promoter in *S. cerevisiae ste12*Δ cells. All strains were co-transformed with a Venus reporter plasmid. Growth arrest (OD_600_) and fluorescence intensity were measured after 20 hours of incubation in the presence or absence of 10 µM peptide. All Data represent the mean ± SD from three biological replicates.

Taken together, these results confirmed previous results that showed that NgSte12(1) could partially substitute for ScSte12 in *S. cerevisiae* and induce mating [19] (in our case, cell cycle arrest). But our results added the new observations that NgSte12(1) functioned constitutively rather than in a pheromone-responsive manner and that simple FUS3 promoter activation did not occur.

The conserved Dig1-binding motif DFPLDYF has been reported in the pheromone-responsive domain (PRD) of Ste12 homologues [37]. Consistent with previous work [19], our multiple sequence alignment showed that NgSte12(1) lacks this motif **(Supplementary Figure 1)**, hinting at the possibility that NgSte12(1) could not be repressed via the inhibitor Dig1.

We thus tested next if any of the NgSte12(1) domains, specifically the PRD with the missing Dig1 motif, might cause the constitutive cell-cycle arrest and/or the inability of FUS3 promoter activation; We thus employed a domain-swapping approach in which each domain of ScSte12 (the DNA-binding domain (DBD), PRD, and transcriptional activation domain (TAD)) was individually replaced by its NgSte12(1) counterpart **(Figure 1A and 2C)**. NgSte12(1) encodes a 599 amino acid protein (versus 689 aa for ScSte12) and the domain boundaries were defined by manually inspecting a multiple sequence alignment of Ste12 homologues from five yeast species **(Figure 2C; Supplementary Figure 1)**. This resulted in defining the DBD as stretching from aa 1–218, the PRD stretching from aa 270–357 and the TAD stretching from aa 410–599. Noteworthy, the DBD of NgSte12(1) shares 65–70% primary sequence identity with homologues across species and was structurally highly conserved, as evidenced by near-complete structural overlap with the ScDBD in a predicted 3D structure (RMSD = 0.417; **Figure 2D)**. The NgSte12(1) TAD showed no predicted secondary structure and shared little sequence homology with other Ste12 proteins. This is expected, as activation domains are known to be sequence diverse and intrinsically disordered [38].

All three domain-swap chimaeras, NgDBD, NgPRD, and NgTAD **(Supplementary Tables 6 and 7)** were expressed from the ScSte12 promoter from the low-copy vector pRS413 in BY4733 *ste12Δ*. Full-length ScSte12, full-length NgSte12, and an empty vector were used as controls.

All three chimaeras showed both growth arrest and fluorescence activation in a pheromone-dependent manner **(Figure 2E)**, demonstrating that each NgSte12 subdomain was individually functional and compatible with its *S. cerevisiae* counterpart. Even NgPRD remained pheromone-responsive despite lacking the consensus Dig1-binding motif, suggesting that this sequence alone is not critical for Dig1/Dig2-mediated repression.

In addition, another unexpected result was observed: When the full-length NgSte12 was expressed under the ScSte12 promoter - a control we had included, given that chimaeras were driven by the ScSte12 native promoter - peptide-dependent activation of cell-cycle arrest was observed, but with weaker growth arrest compared to the individual domain chimaeras (**Figure 2E**). This contrasted with the constitutive growth arrest observed earlier when NgSte12 was driven by its own promoter (**Figure 2A and B**). It suggested that promoter-derived differences in expression levels might contribute to the constitutive phenotype. Ste12 expression levels were shown before to play a critical role in causing constitutive phenotypes, possibly because elevated Ste12 levels saturate Dig1/Dig2 repression, leaving a free pool of active Ste12 [33, 39, 40].

We tested promoter strength by cloning both promoters to drive a Venus reporter. Comparison, however revealed similar basal expression levels between the two promoters **(Supplementary Figure 6)**, leaving the mechanism for the differential phenotypes unresolved. One hypothesis is that both promoters were subject to different positive feedback dynamics: both promoters contain Ste12 response elements, but differing in spacing **(Supplementary Table 5)**. Since the organisation of Ste12 binding sites has recently been shown to directly influence the onset and strength of Ste12-mediated transcription [41], differences in this feedback architecture may account for the distinct expression levels and phenotypes observed with the two promoters.

Collectively, these results demonstrated that each of the three NgSte12(1) domains could functionally replace its corresponding ScSte12 domain while retaining pheromone-responsive signalling in combination with the remaining ScSte12 domains. Also, full NgSte12 could somewhat function as a peptide-responsive, and as such, GPCR-linked transcription factor, dependent on promoter context, indicating a minimally functional peptide-GPCR/reporter cascade could be established with swapped Ng/Sc parts. Minimally functional, being defined by us as some statistically significant difference in growth that could be observed after peptide induction.

### ScSte12 and ScSte2 expression induce peptide-independent FUS3 promoter activation in both *N glabratus* mating types, indicating potential for peptide-GPCR/Ste12/reporter signalling

Next, we investigated if a functional peptide-GPCR/Ste12/reporter cascade could be established in *N. glabratus*, by using the proven-to-be-functional parts from *S. cerevisiae*, specifically the transcription factor Ste12 and in addition the peptide GPCR Ste2. Plasmid-encoded NgSte12(1) under its native promoter was also tested to see if higher gene dosage of Ste12(1) could establish a differential response when compared to genomic expression.

Therefore, plasmid-encoded ScSte12 and NgSte12(1) were introduced into the *N. glabratus* wildtype and *ste12Δ* strains of both mating types and growth arrest and reporter activation were assessed in the presence and absence of 10 µM peptide **(Figure 3A)**. For α-type strains, we would not expect a peptide-inducible phenotype as the strains predominantly express the Ste3 GPCR, but we were nevertheless interested to test the construct to not overlook unexpected phenotypes. All four strains transformed with empty vectors, were used as controls.

**Fig. 3.**
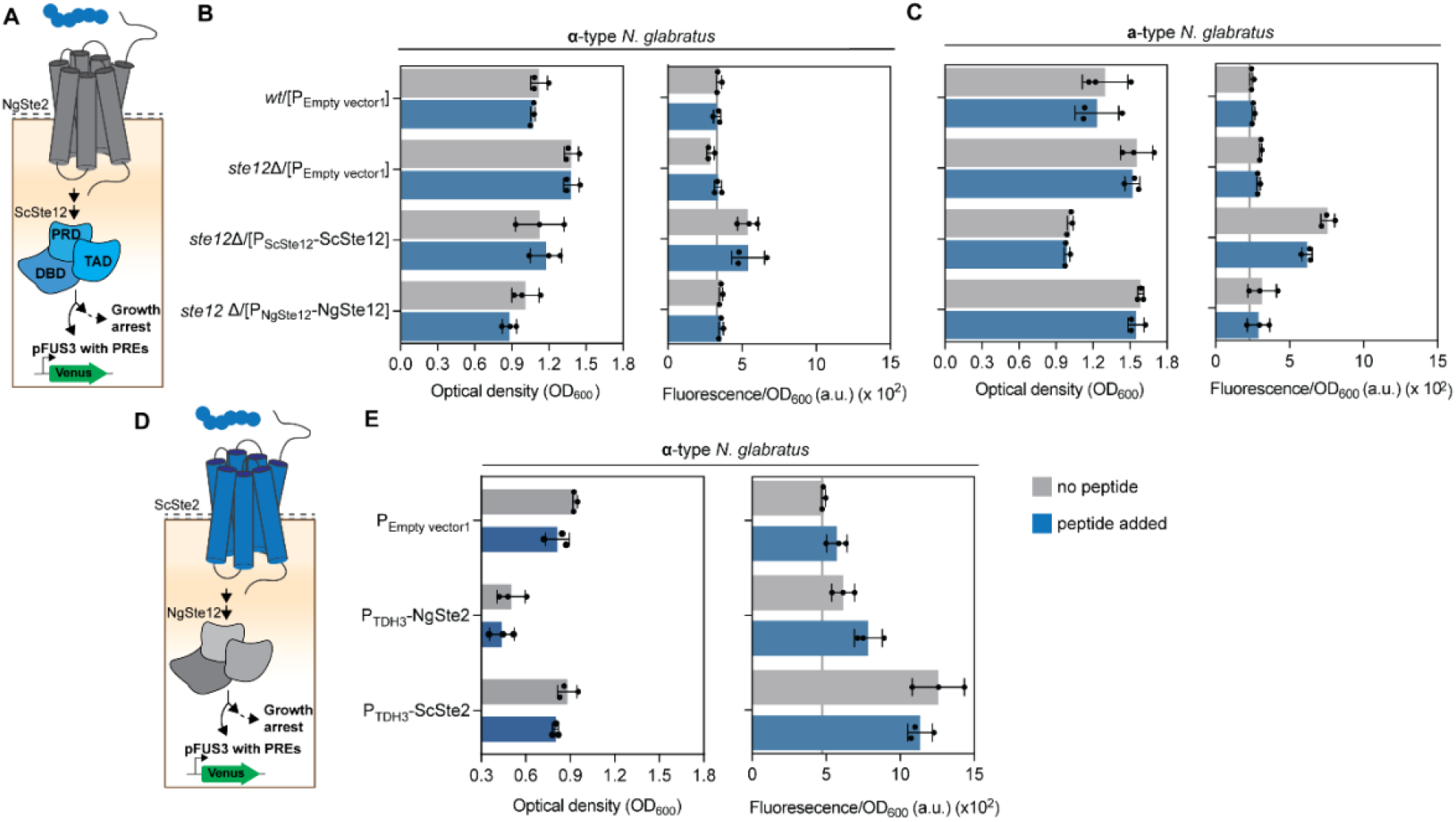
ScSte12 and Ste2 expression in both mating types of *N. glabratus*. **A**. Schematic representation of the chimeric signalling cascade tested to evaluate whether signalling from the endogenous *N. glabratus* peptide–GPCR is functionally transmitted to *S. cerevisiae* ScSte12, thereby inducing FUS3 promoter activation and growth arrest. **B and C**. Functional analysis of ScSte12 and Ngste12 expression in **α-type** (B) and **a-type cells** (C). Ste12 expression plasmids or the Empty vector were used to transform wild-type or *ste12Δ* strains. All strains were co-transformed with a Venus reporter plasmid. Growth arrest and fluorescence intensity were measured when cells were treated with or without 10 µM peptide at 20 h. Data represent the mean ± SD from three biological replicates. **D**. Schematic representation of the chimeric signalling cascade tested to evaluate whether signalling from the heterologous *S. cerevisiae* peptide–GPCR is functionally transmitted to *N. glabratus* NgSte12(1), thereby inducing FUS3 promoter activation and growth arrest. **E**. Ste2 expression plasmids or Empty vector were used to transform wild type *N. glabratus* **α**-type cells. All strains were co-transformed with a reporter plasmid. The growth arrest and fluorescence activation were measure in the presence and absence of 10 µM peptide at 20 h. Data represent the mean ± SD from three biological replicates.

First, none of the *N. glabratus* transformants showed peptide-dependent behaviour **(Figure 3B and C)**. Second, NgSte12(1) deletion and NgSte12(1) plasmid-based expression had no significant phenotype in a-type cells **(Figure 3C)**. In α-type *N. glabratus*, deletion of *ste12*(1) resulted in a slight (17%) decrease in fluorescent levels from the pFUS3 promoter and a slight (23%) increase in final optical density in the absence of 10 µM peptide treatment, which was abolished back to wild-type levels when NgSte2(1) was introduced back, by expression from a plasmid **(Figure 3B and Supplementary Figure 7)**.

Expression of *S. cerevisiae’s* ScSte12, in contrast, caused increased fluorescence when compared with the respective wild-type or *ste12*Δ parent in both mating types, however, independent of peptide (**Figure 3B and C)**. *N. glabratus* a-type cells showed approximately 2.4-fold higher fluorescence **(Figure 3C)**, thus FUS3 promoter activation, and α-type cells showed an approximately 1.6-fold increase (**Figure 3B)**. Elevated activation levels in the absence of peptide might mean that *S. cerevisiae’s* ScSte12 acts as a transcription factor that is insufficiently repressed in *N. glabratus*, either because of too high expression or mechanical lack of repression by Dig1/Dig2 as it would happen in the natural host *S. cerevisiae*. Further, the failure of ScSte12 to respond to pheromone in *N. glabratus*, despite being fully pheromone-responsive in *S. cerevisiae*, may reflect either impaired upstream GPCR-to-Ste12 signalling or elevated basal ScSte12 activity that reduces the dynamic range of the reporter. This GPCR-to-Ste12 signalling impairment could reflect either that the *N. glabratus* peptide-GPCR NgSte2 was not functional or insufficiently expressed, or that it coupled to a different transcription factor or that we were using an incorrect ligand: a point supported by previous findings showing that NgSte2 exhibits very low activation in response to both its own cognate mating peptide and the mating peptide of *S. cerevisiae* when heterologously expressed in *S. cerevisiae* [32].

To test for some of these hypotheses, we expressed ScSte2 in α-type *N. glabratus* wild-type cells (no Ste12 deletion) from the strong TDH3 promoter (**Figure 3D**). *S. cerevisiae’s* Ste2 is a well-characterized functional peptide-GPCR with known ligand, which has shown capacity to couple to NgSte12(1) (**Figure 2E**). We further tested if *N. glabratus*’ peptide-GPCR NgSte2 could lead to a measurable readout when higher expressed (from a TDH3 promoter on a plasmid, rather than only genomically), given that GPCR abundance on the cell surface affects downstream signal readout [46]. An empty vector served as a baseline reference.

Expression of ScSte2 from the strong TDH3 promoter induced fluorescent reporter activation (**Figure 3E**), when compared to the empty vector control; however, in a constitutive fashion, independent of peptide ligand. Still, it showed that ScSte2 could couple to FUS3 activation, constitutively activating the Gα and downstream components.NgSte2 expression from the same strong promoter reduced the final OD_600_ (indicative of growth arrest or metabolic burden, given that it is unclear whether *N. glabratus* has the capacity for this type of cell) and slightly elevated both raw and normalized fluorescence when compared to the empty vector control, may indicate that ectopic NgSte2 expression was insufficient to activate the pathway. It thus appeared that the *S. cerevisiae* components ScSte2 and ScSte12 could establish a signalling link to the FUS3 promoter. For NgSte12(1) in its native signalling context, it remains unclear from which upstream component NgSte12 receives its input. It is also unclear what role the peptide-GPCR NgSte2 plays in *N. glabratus*, whether it senses a different ligand, or if it signals via a different route and, as such, connects to a different transcription factor.

## CONCLUSIONS

Using a component-swapping approach across *S. cerevisiae* and *N. glabratus* (both mating types), we demonstrate that full-length NgSte12(1) and its subdomains are functional and pheromone-responsive transcription factor (components) in *S. cerevisiae*. However, in its native environment, *N. glabratus*’ NgSte12(1) seems decoupled from its cognate upstream GPCR signalling.

An unexpected finding was that the apparent constitutive activity of NgSte12(1) in *S. cerevisiae* that we observed first was not an intrinsic property, but rather a consequence of promoter context, specifically on the promoter of choice. Functionality of NgSte12(1) as a peptide-GPCR-responsive transcription factor was further substantiated by the fact that all three subdomains were functional. This was at first surprising to us, especially for the pheromone response domain (PRD) of NgSte12 (1) as it lacks the Dig1 repression motif DFPLDYF [19] **(Supplementary Figure 1 and 2)**, which in this case does not seem to be critical.

Overall, our findings add to previous studies, which showed that NgSte12(1) could complement a Ste12 deletion in *S. cerevisiae*; however, in these studies, the peptide-dependence or constitutive nature of this phenotype was not directly disentangled [19]. Based on that, our findings support that the failure of NgSte12(1) to support a peptide-GPCR mediated response in *N. glabrata* is not a property of the transcription factor itself, but reflects a disconnect that lies upstream, between the GPCR and the downstream signalling cascade in its native host.

This upstream disconnect is supported by heterologous ScSte2 and ScSte12 expression in *N. glabratus*: Expression of the *S. cerevisiae* peptide-GPCR Ste2 in *N. glabratus* could activate the pathway constitutively, demonstrating that the downstream signalling machinery, including NgSte12(1), is somewhat capable of transducing a GPCR-derived signal. In contrast, plasmid-based expression of the *N. glabratus* GPCR NgSte2 from a strong promoter did not activate signalling, as such gene dosage is not the reason for the unresponsive GPCR/Ste12 cascade.

This suggests that the *N. glabratus* peptide-GPCR NgSte2 either couples to other downstream pathways, or responds to a different yet unidentified ligand, or requires specific cellular or environmental conditions to be active.

Several follow-up questions arise. Identifying whether NgSte2 responds to an alternative ligand or is uncoupled from the downstream G-protein, will be essential for understanding whether and how peptide-GPCR signalling is activated and transmitted downstream. Further, given that components of this pathway, including Ste12, Ste11, Ste20, and Fus3, have been linked to virulence in *N. glabratus* [28], understanding how NgSte12(1) is activated and what gene expression programmes it controls, represents a compelling direction. Finally, the component-swapping framework developed here provides a reasonable starting point for dissecting GPCR signalling pathway functionality in other poorly characterised fungal pathogens.

Finally, our study also provides a genetically tractable and sequenced *N. glabratus* MATa strain (ySB38 HT^−^) next to the already available *N. glabratus* MATα (ATCC2001 HTL^−^), which could be useful for future studies on *N. glabratus* mating types.

## MATERIAL AND METHODS

### Materials

Media were obtained from BD Bioscience (Franklin Lakes, NJ, USA) and Sigma Aldrich (Darmstadt, Germany). Phusion™ High-Fidelity DNA Polymerases was obtained from ThermoFisher Scientific (Bleiswijk, Netherlands). Restriction enzymes were obtained from New England Biolabs (NEB, Leiden, the Netherlands). Synthetic α-factor peptide (≥ 95% purity) was obtained as chemically synthesized peptide from GenScript (Rijswijk, Netherlands). Primers were ordered from Biolegio (Nijmegen, the Netherlands). Sanger sequencing was provided by Macrogen Europe (Amsterdam, the Netherlands). Synthetic DNA (gBlocks) were obtained from Integrated DNA Technologies (IDT, Amsterdam, the Netherlands). Optical density and fluorescence signal measurements were performed in a microplate reader (Tecan, USA). Sterile, transparent, round-bottom 96-well microtiter plates were obtained from Sigma Aldrich (Darmstadt, Germany). Sterile, black, clear-bottom 96-well microtiter plates were obtained from Corning (Corning Inc.). Plasmids and primers used in this study are listed in **Supplementary Table 6 and Table 8**

### Strains and growth media

Yeast strains used in this study are listed in **Supplementary Table 2**. Yeast mutants were obtained by homologous recombination or by using CRISPR/Cas9 as outlined below. Strains were grown in YPD (1% yeast extract, 2% peptone and 2% glucose), in synthetic complete (SC) with appropriate amino acids media, and in SD medium lacking histidine (−His) and /or tryptophan (−Trp) [36]. Nourseothricin was used at a final concentration of 75 µg/mL. Solid media contained 2% agar. Yeast strains were transformed with plasmids using a standard lithium acetate transformation protocol [37]. *E*.*coli* DH5 alpha was used for routine plasmid cloning. *E. coli* was grown in Luria Broth (LB), transformants were selected on kanamycin 50 (µg/mL) and chloramphenicol (50 µg/mL).

### Strain constructions via homologous recombination using the Nourseothricin (Nat) marker and FLP/FRT marker recycling

Except for the ySB38 *ste12* deletion, all gene deletions in BY4733, ATCC2001 HTL^−^ and ySB38 were obtained by homologous recombination using repair fragments encoding a Nourseothricin (Nat) resistance cassette, flanked by ~500 bp homology arms. For the construction of the ySB38 derivative *his3Δ, trp1Δ*, and *ste12Δ* and the BY4733 *far1Δ* strain, the Nourseothricin resistance cassette was amplified with primers encoding two FRT sites and using pYTK078 (from the yeast toolkit [47]) as template DNA. This allowed removal of the Nourseotrhircon cassette after each round of deletion by a flippase. An in-house flippase (FLP) plasmid was cloned by Golden Gate as follows. The FLP fragment was obtained by amplification from plasmid Pv1393 (Addgene)[48] followed by cloning it into vector pRS413 from the CgYTK kit [49] under the control of the HHF1 promoter and the ENO1 terminator from the yeast toolkit [47]. Yeast mutants were cured from the FLP plasmid after each gene deletion round by growth in YPD media. All mutants were verified by colony PCR using primers listed in supplementary Table 8. Strain ATCC2001 HTL^−^ *ste12Δ* was verified by full genome sequencing as outlined below.

### Strain construction via CRISPR-Cas9 to delete *ste12* in ySB38 *his3*Δ::*frt trp1*Δ::*frt*

The MoClo Yeast toolkit [38] and the CgYTK extension [22] were used to construct a CRISPR-Cas9 system for genome editing of ySB38 *his3*Δ::*frt trp1*Δ::*frt*. The Cas9 gene (pYTK036) was cloned into a pRS413-type vector that was assembled from the YTK components, with a TEF2 promoter (pYTK014) and ENO1 terminator (pYTK051). Chopchop (https://chopchop.cbu.uib.no/) was used to design the sgRNA for NgSte12(1) deletion in *N. glabratus* ySB38 (gRNA26-F and gRNA26-R, **Supplementary Table 8**). A single gRNA was cloned into pYTK050 and the gRNA expression cassette was then cloned into pRS414 (which was assembled from the CgYTK components). Thermocycling conditions for Golden Gate reactions were as follows: 42 °C for 2 min, 16 °C for 5 min, 25 cycles, then 60 °C for 10 min and 80 °C for 10 min.

To construct ySB38 *ste12Δ*, the Cas9 and gRNA expression plasmids were than co-transformed together with the Nourseothricin resistance cassette repair fragment (generated as outlined above with 500 bp homology arms) into ySB38 *his3Δ::frt trp1Δ::frt*.

### Full genome sequencing of ATCC2001 HTL^−^ *ste12Δ* and ySB38 *his3Δ::frt trp1Δ::frt ste12Δ*

Genomic DNA was extracted using the MasterPure Yeast DNA Purification Kit (LGC Biosearch Technologies). The concentration of genomic DNA was measured with Quant-iT™ PicoGreen™ dsDNA Assay Kits and dsDNA Reagents (ThermoFisher Scientific) and then sent for Illumina-based sequencing. Read quality was assessed using FASTQC, subsequently, Illumina reads were trimmed using trimmomatic SE “SLIDINGWINDOW:30:28 ILLUMINACLIP:NexteraPE-PE.fa:2:30:10” [50, 51]. To verify the deletion of *his3, trp1* and *ste12*, we used a two-fold approach. First Illumina reads were mapped to the Ste12, His3 and Trp1 ORFs and the *N. glabratus* ySB38 genome sequence using a custom Python script that calls bowtie2 and processes reads using SAMTOOLS [52, 53]. Second, the Illumina reads were used to assemble a genome using SPADES with default settings. Scaffolds with a coverage below 10 were filtered using a custom Python script. Illumina reads were mapped to the assembly, and duplicates were removed using picard MarkDuplicates REMOVE_DUPLICATES TRUE [54]. Reads were subsequently realigned using gatk3.8 IndelRealigner [55]. The realigned reads were used to polish the genome using Pilon at mindepth 20 [56]. The polished genome was evaluated using QUAST using the *N. glabratus* CBS138 genome published on the *N. glabratus* genome database and deposited on Zenodo (DOI: 10.5281/zenodo.21838110) (20260807) as reference. Genome completeness was assessed through BUSCO using the saccharomycetes_odb10 database [57]. Bedgraph files were created using bedtools GENOMECOV and tracks were visualized using IGV [58, 59]. After verifying genome completeness, the *ste12*, FRT scar (GAAGTTCCTATTCtctagaaaGAATAGGAACTTC) and the *his3* and *trp1* up and downstream regions were aligned to the newly assembled genome (**Supplementary Table 4** and **Figure 4**).

### Construction of Ste12 expression vectors and Venus reporter plasmids

All Ste12 expression plasmids were constructed using Gibson assembly and using the low copy vector pRS413 for expression. pRS413 from the CgYTK was digested with BsaI to release the GFP dropout cassette, followed by treatment with alkaline phosphatase to remove 5’- and 3’-phosphate groups.

The ScSte12 and NgSte12(1) genes were amplified from the genomes of *S. cerevisiae* BY4741 and *N. glabratus* ATCC2001HTL^−^, respectively, along with their individual promoters. Terminator ENO1 was amplified by PCR using the primer pair in **Supplementary Table 8**.

All Ste12 chimeras were driven by the ScSte12 promoter and terminated by the ENO1 terminator. To generate the Ste12 chimeras’ individual domains were amplified with primer pairs listed in **Supplementary Table 8**. These fragments were then assembled into the pRS413 vector, to construct the final plasmids. To assess the activity of ScSte12, NgSte12(1) and chimeric Ste12, the Venus reporter plasmid was constructed. NgFus3 promoter (pSB95, CgYTK), Venus reporter (pYTK033), and ENO1 terminator (pYTK051) were assembled into the pRS414 vector using Golden Gate cloning.

To compare the strengths of ScSte12 and NgSte12(1) promoter, two Venus expression plasmids, PScSte12-Venus and PNgSte12(1)-Venus, were constructed using the pRS413 vector. For the PScSte12-Venus construct, Venus (pYTK033) and the ENO1 terminator (pYTK051) were first assembled by Golden Gate cloning to generate a linear Venus-ENO1 fragment. This fragment was then assembled with the PCR-amplified ScSte12 promoter using Gibson Assembly. For the PNgSte12(1)-Venus construct, the NgSte12(1) promoter was synthesised as a gBlock by GenScript and assembled with Venus and the ENO1 terminator before being cloned into the pRS413 vector.

### Growth arrest and fluorescence reporter assays

The assays were performed as previously reported [60]. In brief, strains were grown overnight in 2 mL of SD-His/Trp and the resulting overnight culture was adjusted to an approximate OD_600_ of 3 (measured in a cuvette-based spectrophotometer). This suspension served as a concentrated cell suspension to set up the 96-well assay. The 96-well assay was set up in transparent, round-bottom 96-well microtiter plates using a 200 µL total volume. The following components were mixed: 100 µL 2x SD-His/-Trp, 5 µL cell suspension, 20 µL of 100 µM peptide solution and 75 µL sterile H_2_O. Cells were incubated at 30 °C for 20 h and subsequently transferred to a black, clear-bottom 96-well microtiter plate for measurement of OD_600_ and fluorescence using a microplate reader (Tecan, USA). Fluorescence was measured with an excitation wavelength of 488nm and an emission wavelength of 530nm. The OD_600_ value was corrected as described before to account for detector saturation [60]. The reported fluorescence intensity/OD_600_ was calculated by dividing the fluorescence intensity by the normalized OD_600_. All experiments were performed with three biological replicates, and data are presented as the mean ± standard deviation (SD).

### Statistical analysis and definition of replicates

All experiments were performed in three independent biological replicates. Each biological replicate was performed using an independently grown overnight culture. Data are presented as the mean ± SD. Statistical significance was determined using Student’s t-test.

## Contributions

**M.L**.: Conceptualization, Investigation, Formal analysis, Data curation, Writing - original draft.

**T.A.D.V**.: Formal analysis, Data curation, Writing - review & editing.

**S.B**.: Conceptualization, Supervision, Data curation, Writing – original draft and review & editing.

## Data availability

The genome sequencing data have been deposited in Zenodo (DOI: 10.5281/zenodo.21838110).

