## Supplementary Information for "The transcription factor Ste12(1) from *Nakaseomyces glabratus* acts pheromone-responsive in *S. cerevisiae* but seems decoupled from its cognate upstream GPCR in its native host"

**SUPPLEMENTARY TABLES**

**Supplementary Table 1. Homologs of *Saccharomyces cerevisiae* pheromone response pathway components in *Nakaseomyces glabratus*.** % query coverage and % amino acid sequence identity is given in comparison to the *Saccharomyces cerevisiae* components.

| Protein function | *S. cerevisiae* | *N. glabratus* | Query coverage | Amino acid sequence identity [a] |
|---|---|---|---|---|
| GPCR | Ste2 | Ste2 (CAGL0K12430g) | 96% | 43% |
| G-protein | Gpa1 | Gpa1 (CAGL0F06677g) | 100% | 70% |
| | Ste4 | CAGL0L02761g | 97% | 57% |
| | Ste18 | CAGL0M09207g | 83% | 60% |
| MAPKKK | Ste11 | Ste11 (CAGL0B02739g) | 96% | 64% |
| MAPKK | Ste7 | CAGL0I03498g | 85% | 55% |
| MAPK | Fus3 | Fus3 (CAGL0J04290g) | 100% | 79% |
| Transcription factor | Ste12 | Ste12(1) (CAGL0M01254g) [b] | 36% | 67% |
| | | Ste12(2) (CAGL0H02145g) | 25% | 64% |
| Repressor | Dig1 | Dig1 (CAGL0L12782g) | 23% | 43% |
| | Dig2 | Not found [c] | | |

[a] identify of amino acid sequences was analysed using BLASTP.
[b] used in this study.
[c] Searching for ScDig2 homolog in *N. glabratus* using the Candida Genome Database.

**Supplementary Table 2. Strains were used in this study**

| **Strain name** | | **Genotype** | **Reference** |
|---|---|---|---|
| BY4741 | Strain 2 | MATa *his3Δ1 leu2Δ0 met15Δ0 ura3Δ0* | [1] |
| BY4733 *ste12Δ::meth1* | Strain 1 | MATa *his3Δ200 leu2Δ0 met15Δ0 trp1Δ63 ura3Δ0 ste12Δ::meth1* | [2] |
| ATCC2001 HTL- | Strain 5 | MATα HTL- | [3] |
| BY4733 | | MATa *his3Δ200 leu2Δ0 met15Δ0 trp1Δ63 ura3Δ0 ste12Δ::meth1 far1Δ::nat* | This study |
| ATCC2001 HTL- *ste12Δ:: nat* | Strain 3 | MATα HTL- *ste12Δ::nat* | This study |
| ySB38 | | MATa | Billerbeck Lab, gift from Dr. Daniel Green, Columbia University Medical Center |
| ySB38 *his3Δ* | | MATa *his3Δ::frt* | This study |
| ySB38 *his3Δ::frt trp1Δ::frt* | Strain 6 | MATa *his3Δ::frt trp1Δ::frt* | This study |
| ySB38 *his3Δ::frt trp1Δ::frt ste12Δ::nat* | Strain 4 | MATa *his3Δ::frt trp1Δ::frt ste12Δ::nat* | This study |

**Supplementary Table 3. mRNA read counts for NgSte2, NgSte3, NgSte12(1) and NgSte12(2) in strain *N. glabratus* ATCC2001.** Data were extracted from [2], Supplementary Table 1. Actin and triosephosphate isomerase were chosen as house-keeping genes, for comparison.

| **ORF ID** | **Protein name** | **Mean RPKM (Reads per kilobase of transcript per million mapped reads** | |
|---|---|---|---|
| | | **GSNO** | **YPD** |
| CAGL0K12430g | Ste2 | 0.358767019 | 0.584983759 |
| CAGL0M08184g | Ste3 | 14.8656017 | 46.99634306 |
| CAGL0M01254g | Ste12(1) | 13.90358477 | 7.332586004 |
| CAGL0H02145g | Ste12(2) | 9.47868092 | 8.357672735 |
| CAGL0K12694g | Act1 (Actin) | 796.7580262 | 1433.180641 |
| CAGL0H08327g | Tpi (Triose-phosphate isomerase) | 342.6878784 | 1285.044506 |

**Supplementary Table 4. Verification of *trp1* and *his3* deletion from ySB38 genome by full genome sequencing (Illumina-based)**. Mapping positions of the FRT site relative to 999 bp downstream regions of *his3* and *trp1*

| **ORF** | **Contig** | **Mapping position FRT scar** | **Downstream** |
|---|---|---|---|
| *trp1* | NODE200 | 446-479 | 480-1479 |
| *his3* | NODE207 | 3215-3248 | 3248-4247 |

**Supplementary Table 5. Promoter sequences of ScSte12 and NgSte12 used in this study.**
Nucleotides highlighted in yellow are pheromone response elements (PREs) or PRE-like sites.

| Promoter | Sequence (5'-3') |
|---|---|
| ScSte12 | CTTACTTACATCTGAAAATTGCAAGTTACATTCTTTGTATAACGAACGTTA<br>AGGAACCCATAAAGCTAAAGACATTTGTTGAAAACGAATGTAAAGAATTG<br>GTCCAGTTTGCACAAGACACCCTGAAGAACTTCGTTCAGTAATCACTTTCA<br>AGCTGTAGTATGTAAACGATATAGATGAAGTTTTCGTGTGTATAAATATAT<br>GAACTCTAGAGTGTTGCATAATTTGAAACACAGCATTTCTTTTCGGAGAGC<br>TCGTTTCAAAATGAAACAAACGCCGTTGTCCGTTTTCGTCTCAATAGAAAA<br>AGTGAAACAGATAAAAATTGTTTTAAAAGAAACGAATTTGCAACATCTTA<br>AGATATATCAAAACTAATACAAAACAGCCTAAAAAAGATTGAACAACTCT<br>TCGCGGTCAGGTCTCGACACCATAAATCGAAGTACTCGTACGCTAGTTTTC<br>TCGCACATAGTACCACTACGTTCCTTTTACAATTAGATTACTTCTTTTTAGT<br>TGACTTTTTTGAGACGTTCGTGCCATTCATAAAATAGGAAAAGATAACAGG<br>TAAGCACTGAAGACTTGTTTTATAAGTGTCCCAAGCGAGACCTAGAGTGG<br>ATATTGATATTTCTCAAACAAGACTCGTCGAAGAAAACACACTTTTATAGC<br>GGAACCGCTTTCTTTATTTGAATTGTCTTGTTCACCAAGG |
| NgSte12(1) | GGCAATTCGTCGCAATACTACAAATTCTCTCTACTGGAAAATAAGGTTTTG<br>AAACTGTTTACTACTGTCAATGATAAAAAACTTCCTTACACCAGTCTTCAT<br>GAACTAAAATTAAAAATCATGAAACTTAATATAGTTTAACTGTTACCAAGG<br>AACACTAAATCTGTCCTAATTTTAAGACCACTGACAGATTCTTTTCATTTTA<br>CCTACTATCTAGACCCTCCTGACTCAGAACCCTGGCATTGTTTTTTGTTTTA<br>CTTGTTTAAAACTGATATGAAACATCCATGTTACAAAATGGGATGGTAATG<br>GAACCATTTGATATTTCATTATAAAACATAGTGTCTTGACTTCAGTTTTTGT<br>TTTGATCCTAATTCTTTGATAATTTTGACTCAGGACTGTAACAGATAAACG<br>AGGATTTATTATTCTGATTTTTTTTTGTTGACATATTCTGAGTTAGATCAAT<br>TGAGTGGTTTTCCAAGACAAATCACATATACCAAATAAAAATATCATTATT<br>GATCACCCAGTTGCAAACATCTGATATATTGATAACTCCTTTGATAACGTG<br>TTTGTTTATTGATAGTTCATAGGAATTGAACACTAATAAAATTTGTTGGTTC<br>TGCCAAAAAATAGCGAAAACCGGTAGTCTATTTCCACGCCGATAATTAAT<br>ATTATACCAAATATACTACAACGAGAAGTTATATTCTGTAATATCTACACC |

**Supplementary Table 6. Plasmids used in this study.**

| Plasmid | Construct | Source | Description |
|---|---|---|---|
| pRS413 | Vector Vs1.1 | [4] | Plasmid vector |
| pRS414 | Vector Vs2.2 | [4] | Plasmid vector |
| pML001 | pTEF2-Cas9-tENO1-pRS413 | This study | Ste12 deletion |
| pML68 | gRNA26-pYTK050 | This study | |
| pML73 | gRNA26-pRS414 | This study | |
| Pv1393 | Sap2-FLP | Addgene #111430 [5] | NatR marker recycling |
| pML85 | pYTK001-FLP | This study | |
| pML86 | HHF1-FLP-tENO1-pRS413 | This study | |
| ScSte12(pML204) | ScSte12p-ScSte12-tENO1-pRS413 | This study | Ste12 and variants |
| NgSte12(pML84) | NgSte12p-NgSte12-tENO1-pRS413 | This study | |
| ScSte12 (pML106) | pScSte12-ScSte12p-alfa-tENO1-pRS413 | This study | |
| NgDBD (pML107) | pScSte12-NgDBD-alfa-tENO1-pRS413 | This study | |
| NgPRD (pML110) | pScSte12-NgPRD-alfa-tENO1-pRS413 | This study | |
| NgTAD (pML111) | pScSte12-NgTAD-alfa-tENO1-pRS413 | This study | |
| NgSte12 (pML113) | pScSte12-NgSte12-alfa-tENO1-pRS413 | This study | |
| pML147 | NgSte12p-Venus-teno1-tENO1-pRS413 | This study | Ability of Ste12 promoter |
| pML148 | ScSte12p-Venus-tENO1-pRS413 | This study | |
| ScSte2(pML173) | TDH3-ScSte2-tENO1-pRS413 | This study | Ste2 plasmid |
| NgSte2(pML174) | TDH3-NgSte2-tENO1-pRS413 | This study | |
| Empty vector1 (pML103) | pRS413-empty vector | This study | Negative control of Ste12 and Ste2 plasmids |
| pSB95 | pYTK001-NgFUS3p | This study | Reporter plasmid |
| pML83 | NgFus3-Venus- tENO1-pRS414 | This study | |
| Empty vector2 (pML76) | pRS414-empty vector | This study | Negative control of Venus reporter |

**Supplementary Table 7. Amino acid sequences of Ste12 chimeras.** Amino acids highlighted in yellow are derived from the indicated NgSte12(1) domain, amino acids highlighted in blue indicate ALFA tag sequence, whereas all other residues remain those of ScSte12.

| Ste12 variants | Amino acid sequences |
|---|---|
| NgDBD (pML107) | MSPFIKGRQRRTEEILKGDLKHHNDDGLVKSVIDDLEDAISAIEDLKFFLA<br>TAPLNWHENQVIRRYYLNNSQGFISCVFWNSLYYMTGTDIVKACMYRM<br>EKFGRKVIERKKFEEGLFSDLRNLKCGLDATLEQPKSKFLKFLFRNLCLKT<br>QKKQKVFFWFSIPHDKLFADALERDLKREFNGQNPTTIAIQEPALSFNYD<br>QDSKLSLQEQLSRHISTKRPSSTTKSDNSPPKLESENFKDNELVTVTNQPL<br>LGVGLMDDDAPESPSQINDFIPQKLIIEPNTLELNGLTEETPHDLPKNTAKG<br>RDEEDFPLDYFPVSVEYPTEENAFDPFPPQAFTPAAPSMPISYDNVNERDS<br>MPVNSLLNRYPYQLSVAPTFPVPPSSSRQHFMTNRDFYSSNNNKEKLVSP<br>SDPTSYMKYDEPVMDFDESRPNENCTNAKSHNSGQQTKQHQLYSNNFQ<br>QSYPNGMVPGYYPKMPYNPMGGDPLLDQAFYGADDFFFPPEGCDNNML<br>YPQTATSWNVLPPQAMQPAPTYVGRPYTPNYRSTPGSAMFPYMQSSNSM<br>QWNTAVSPYSSRAPSTTAKNYPPSTFYSQNINQYPRRRTVGMKSSQGNVP<br>TGNKQSVGKSAKISKPLHIKTSAYQKQYKINLETKARPSAGDEDSAHPDK<br>NKEISMPTPDSNTLVVQSEEGGAHSLEVDTNRRSDKNLPDATPSRLEEEL<br>RRRLTEP |
| NgPRD (pML110) | MKVQITNSRTEEILKVQANNENDEVSKATPGEVEESLRLIGDLKFFLATAP<br>VNWQENQIIRRYYLNSGQGFVSCVFWNNLYYITGTDIVKCCLYRMQKFG<br>REVVQKKKFEEGIFSDLRNLKCGIDATLEQPKSEFLSFLFRNMCLKTQKK<br>QKVFFWFSVAHDKLFADALERDLKRESLNQPSTTKPVNEPALSFSYDSSS<br>DKPLYDQLLQHLDSRRPSSTTKSDNSPPKLESENFKDNELVTVTNQPLLG<br>VGSKFQSCNTSHAQSAISSPLSPTNNLPASMDTDYSSDLKELTSDSPNFDN<br>DDNFMNIKIEYPDENISNGFLATSLNPDQSFLYYDDRNMEIDKNSSIDTNS<br>LLNRYPYQLSVAPTFPVPPSSSRQHFMTNRDFYSSNNNKEKLVSPSDPTSY<br>MKYDEPVMDFDESRPNENCTNAKSHNSGQQTKQHQLYSNNFQQSYPNG<br>MVPGYYPKMPYNPMGGDPLLDQAFYGADDFFFPPEGCDNNMLYPQTAT<br>SWNVLPPQAMQPAPTYVGRPYTPNYRSTPGSAMFPYMQSSNSMQWNTA<br>VSPYSSRAPSTTAKNYPPSTFYSQNINQYPRRRTVGMKSSQGNVPTGNKQ<br>SVGKSAKISKPLHIKTSAYQKQYKINLETKARPSAGDEDSAHPDKNKEIS<br>MPTPDSNTLVVQSEEGGAHSLEVDTNRRSDKNLPDATPSRLEEELRRRLT<br>EP |
| NgTAD (pML111) | MKVQITNSRTEEILKVQANNENDEVSKATPGEVEESLRLIGDLKFFLATAP<br>VNWQENQIIRRYYLNSGQGFVSCVFWNNLYYITGTDIVKCCLYRMQKFG<br>REVVQKKKFEEGIFSDLRNLKCGIDATLEQPKSEFLSFLFRNMCLKTQKK<br>QKVFFWFSVAHDKLFADALERDLKRESLNQPSTTKPVNEPALSFSYDSSS<br>DKPLYDQLLQHLDSRRPSSTTKSDNSPPKLESENFKDNELVTVTNQPLLG<br>VGLMDDDAPESPSQINDFIPQKLIIEPNTLELNGLTEETPHDLPKNTAKGR<br>DEEDFPLDYFPVSVEYPTEENAFDPFPPQAFTPAAPSMPISYDNVNERDSM<br>PVNSLLNRYPYQLSVAPTFPVPPSSSRQHFMTPFQDIKSAAKDGAFTTQKI<br>NPSVGNNLHFPYPEVTSPGLVDSITLSELLDENCNNASFELNNERVKISSQ<br>ELSIWNMLPHNSTSHMTPVFHSLLSAGIKGTHLPAYSPFLTTTPWGSLSSP<br>FSSAKSNKSSQRRQSQFDTRTLGKRRKTKTCSKYSSPNGIARNTKPNKVL<br>KNSINKMIDTLRSNTKNVSPSRLEEELRRRLTEP |

**Supplementary Table 8. Primers used in this study.**

| Primer | Primer sequence (5'-3') | Application |
|---|---|---|
| ML109 | GACTTTTAGCGACAGCACCACTTAAT | gRNA used for *ste12* deletion in *N. glabratus* ySB38 |
| ML110 | aaacATTAAGTGGTGCTGTCGCTAaa | |
| ML104 | TATACTACAACGAGAAGTTATATTCTGTAATATCTACACCtacagtttagcttgcctcgt | *ste12* deletion in *N. glabratus* ATCC2001H $TL^-$ |
| ML105 | aattatctATAAATTGATCCTTTTAAATTAACATGATATCactccagtatagcgaccagc | |
| ML106 | GTTAGATCAATTGAGTGGTT | *ste12* deletion verification in *N. glabratus* ATCC2001H $TL^-$ |
| ML107 | AGCAGTTATCATTACAGGAATA | |
| ML108 | tcttctataagcatggatagca | |
| ML53 | tacagtttagcttgcctcgt | Donor DNA (NatR cassette) for genes deletion |
| ML54 | cagtatagcgaccagcattc | |
| Ml138 | GAGCTCGATTCCCCCATG | *his3* deletion |
| ML139 | cggcggggacgaggcaagctaaactgtaGAAGTTCCTATTCtttctagaGAATAGGAACTTCCCTTTGCTCGATGCTTCTCT | |
| ML140 | atgtgaatgctggtcgctatactgGAAGTTCCTATTCtctagaaaGAATAGGAACTTCAACACAGCCCACAGCTACCACC | |
| ML141 | TGCTCTGCTAACTCAGTCAT | |
| ML142 | TCGGTGCTCTACAGGAATC | *trp1* deletion |
| ML143 | ggcggggacgaggcaagctaaactgtaGAAGTTCCTATTCtttctagaGAATAGGAACTTCCTTGTTGGTGTGTTGGTTATGT | |
| ML144 | tgtgaatgctggtcgctatactgGAAGTTCCTATTCtctagaaaGAATAGGAACTTCATTACGAAGAGAAGTCTCAATGAGG | |
| ML145 | ACTAAATTTGGTGAAGATACTTCTGTGG | |
| ML177 | CCCTCCTGACTCAGAACCCT | *ste12* deletion and verification in ySB38 $HT^-$ |
| ML178 | acgaggcaagctaaactgtaGAAGTTCCTATTCtttctagaGAATAGGAACTTCGGTGTAGATATTACAGAATATAACTTC | |
| ML181 | tgaatgctggtcgctatactgGAAGTTCCTATTCtctagaaaGAATAGGAACTTCGATATCATGTTAATTTAAAAGGATCAATT | |
| ML182 | TGCAATTTTATCCGAAGAGTC | |
| ML165 | TACAAATTCTCTCTACTGGAAAA | |
| ML166 | GACGAGGTTTCAACATCATTG | |
| ML275 | TATAGGTGATATCCTGGAGAGAC | *far1* deletion and verification in BY4733 *ste12Δ::meth1* |
| ML276 | acgaggcaagctaaactgtaGAAGTTCCTATTCtttctagaGAATAGGAACTTCTAATAGATTGCCTTCTTACGCCCACG | |
| ML277 | atgtgaatgctggtcgctatactgGAAGTTCCTATTCtctagaaaGAATAGGAACTTCTAGTTCGGGAATCGAGGCCCGTATTTC | |
| ML278 | GTGGAAATCGTATGGAATG | |
| ML279 | CAAAATTAGTTTATTGGGCC | |
| ML54 | cagtatagcgaccagcattc | |
| ML146 | GCATCGTCTCATCGGTCTCATATGccacaatttgatatattatg | Construction of FLP plasmid for marker recycling |
| ML147 | ATGCCGTCTCAGGTCTCAGGATttatatgcgtctatttatgtagg | |

| | | |
|---|---|---|
| ML131 | agtacagacactgcgacaacgtggcaattcgtcgcaatacCTTACTTACATCTGAAAATTGCAAGTTACATTCTTTG | constructions of Ste12 expression plasmids |
| ML132 | aaaagctctcgagttaggatTCAGGTTGCATCTGGAAGGTTTTTATCG | |
| ML133 | ACCTTCCAGATGCAACCTGAatcctaactcgagagcttttgattaagc | |
| ML134 | catgattcgtcagtgtgagcaccactgacgagcagatttcatacatgggtgaccaaaagagcg | |
| ML127 | agtacagacactgcgacaacgtggcaattcgtcgcaatacTACAAATTCTCTCTACTGGAAAATAAGGTTTGAAACTGTTTACTACTGt | |
| ML128 | aaaagctctcgagttaggatCTAACTTacatttttgtatttgaaCGTAAGGTATCTATCATTTTG | |
| ML129 | atacaaaaaatgtAAGTTAGatcctaactcgagagcttttgattaagc | |
| ML131 | agtacagacactgcgacaacgtggcaattcgtcgcaatacCTTACTTACATCTGAAAATTGCAAGTTACATTCTTTG | Constructions of Ste12 chimeras |
| ML187 | AATTCTTCTTCCAATCTAGATGGGGTTGCATCTGGAAGG | |
| ML186 | CCATCTAGATTGGAAGAAGAATTGAGAAGAAGATTGACTGAACCAtagatcctaactcgagagcttttg | |
| ML48 | tgattcgtcagtgtgagcaccactgacgagcagatttcatacatgggtgaccaaaaga | |
| ML241 | ctACCCTTTATAAATGGGCTCATCCTTGGTGAACAAGACAATTCAAATAAAG | |
| ML189 | ATGAGCCCATTTATAAAGGGTag | |
| ML190 | CTTTGTCGAAATATGGCGGCT | |
| ML191 | GCCGCCATATTTCGACAAAGAGACCATCTAGTACAACAA | |
| ML219 | GTGTTACagctttgaaattttgaGCCAACGCCTAAAAGCG | |
| ML220 | tcaaaatttcaaagctGTAACAC | |
| ML221 | GGGTATCTATTAAGAAGAGAATTTGTATCAATGCTGGAAT | |
| ML222 | AATTCTCTTCTTAATAGATACCCC | |
| ML223 | TTAATATCCTGAAATGGTGTCATAAAATGTTGCCTCGATG | |
| ML224 | ACACCATTTCAGGATATTAAATCAGCC | |
| ML198 | TCTTCTTCCAATCTAGATGGACTTacatttttgtat | |
| ML188 | ctACCCTTTATAAATGGGCTCATCCTTGGTGAACAAGACA | |
| ML273 | attcgtcgcaatacAACGCTTACTTACATCTGAAAATTGC | Construction of ScSte12p-Venus plasmid |
| ML274 | taattcttcacctttagacataCCTTGGTGAACAAGACA | |
| ML269 | GGTCTCAAACGTACAAATTCTCTCTACTGGAAAA | Construction of NgSte12p-Venus plasmid |
| ML270 | GGTCTCACATAGGTGTAGATATTACAGAATA | |

### SUPPLEMENTARY FIGURES

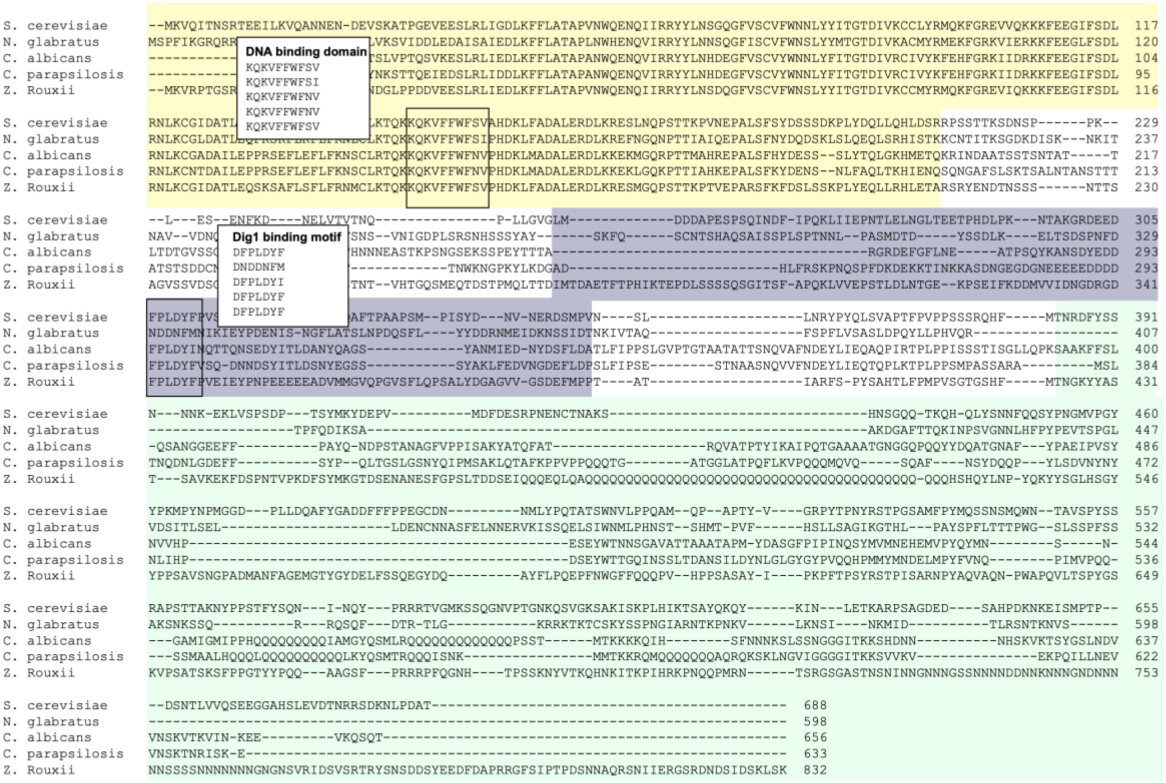


**Supplementary Figure 1. Multiple sequence alignment of Ste12-type transcription factors** from *S. cerevisiae*, *N. glabratus* (Ste12(1)), *Candida albicans*, *Candida parapsilosis*, and *Zygosaccharomyces rouxii*. Alignment was performed using Clustal Omega. Regions highlighted in yellow, purple, and cyan indicate the DNA-binding domain (DBD), pheromone-responsive domain (PRD), and transcriptional activation domain (TAD). The conserved DNA-binding motif and Dig1-binding motif are indicated in the alignment. Notably, the Dig1-binding motif is absent from the NgSte12(1) protein in *N. glabratus*.

```
ScSte12       --MKVQITNSRTEEILKVQANN-ENDEVSKATPGEVEESLRLIGDLKFFLATAPVNWQENQIIRRYYLNSGQGFVSCVFWNNLYYITGTDIVKCCLYRMQKFGREVVQKKKFEEGIFSDL  117
NgSte12(1)    MSPFIKGRQRRTEEILKGDLKHHNDDGLVKSVIDDLEDAISAIEDLKFFLATAPLNWHENQVIRRYYLNNSQGFISCVFWNSLYYMTGTDIVKACMYRMEKFGRKVIERKKFEEGLFSDL  120
NgSte12(2)    --------------------------MMASDDRAVIGNALKSIDELTYFLATAPVNWQPGQVIRRYFLNEELGYISCVFWDNIFYITGTDIIKICIYKMLLFGRVVLQKKKFEEGIFSDL  94
                                          DNA-binding motif
ScSte12       RNLKCGIDATLEQPKSEFLSFLFRNMCLKTQKKQKVFFWFSVAHDKLFADALERDLKRESLNQ--PSTTKPVNEPALSFSYDSSSDKPLYDQLLQHLDSRRPSSTTKSDNSPPK---LES  232
NgSte12(1)    RNLKCGLDATLEQPKSKFLKFLFRNLCLKTQKKQKVFFWFSIPHDKLFADALERDLKREFNGQ--NPTTIAIQEPALSFNYDQDSKLSLQEQLSRHISTKKCNTITKSGDKDISKNKITN  238
NgSte12(2)    RCLKIGEHASLEQPKSPFLDFLYKNMCVKTQKKQKVFYWFSVPHDKLISDALERDLKRESNGMNQLITTRAVREPALTFKYNRSIAGSVFQQARNYFVLRSRIIFSDVFSTPIDVNRAPN  214
                                                                                                         Dig1-binding motif
ScSte12       E-----NFK---DNE---LVTVTNQPLLGVGLMDDDAPESPSQINDFI------PQKLIIEPNTLELNGLT----EETPHDLPKNTAKGRDEEDFPLDYFPVSVEYP-----TEENAFDP  326
NgSte12(1)    AVVDNQNEEQKEANDISSLEEVSNSVNIGDPLSRSNHSS-SYAYSKFQSCNTSHAQSAISSPLS-PTNNLPASMDTDYSSDLKELTS-DSPNFDNDDNFMNIKIEYP-----DENISN-G  349
NgSte12(2)    S-----NQ-----------RVTGNPNDLGVNLSSSQDTVETS-SQKYAASNQSY--SSVHTPI--PSSDTTP---HTQHADLRNMKP----SNDR---AVSKTSQVPPKDNETVNISKQQ  303

ScSte12       FPPQAFTPAAPSMPISYDNVNERDSMPVNSLLNRYPYQLSVAPTFPVPPSSSRQHFMTNRDFYSSNNNKEKLVSPSDPTSYMKYDEPVMDFDESRPN-----------------------  423
NgSte12(1)    FLATSLNP---------------------------------------------------------------------DQSFLYYDDRNMEIDKNSSIDTNKIVTAQFSPFLVSASLDPQY  400
NgSte12(2)    FMPALIDPTTG--------------------------------------------FFEGVQYYISNMGSKEYMCKQRNTPAVN---LNADIKQSSPDNL------------IEKEEDPHA  364

ScSte12       -----------ENCTNAKSHNSGQQTKQHQLYSNNFQQSYPNGMVPGYYPKMPYNPMGG-------------------------------------------------------------  471
NgSte12(1)    LLPHVQRTPF-QDIKSAAKDGAFTTQKINPSVGNNLHFPYPEVTSPGLVDSITLSEL---------------------------------------------------------------  456
NgSte12(2)    SIQ-V-EGPLRCSCGY-SARNTGRSTASSSSSDNSALQSTDDGSDPGILQSRPPPPPSPPPPQFKPEQDPDRGSPTNSASRMGTQVEQKDSFHEYTTSSNEETIRSGVSVAHTQRSGSQF  481

ScSte12       -DPLLDQAFYGADDFFFPPE--GCDN---------NMLYPQTATSWNVLPPQAMQPAPTY---VGRPYTPNYRSTPGSAMFPYMQSSNSMQWNTAVSPYSSRAPSTTAKN----------  566
NgSte12(1)    ----L-------------DE--NCNNASFELNNERVKISSQELSIWNMLPHNSTSHMTPV---FHSLLSAGI---KGTHLPAYSPFLTTTPWGSLSSPFSSAKSNKSSQR----------  541
NgSte12(2)    DDGLASAETYDKDDYNIAGRTSGSDHDDYDNHS-------IESALDIVLPAGTEGNTSSMISTDDEFYSTFMEPFDAS--KYFLSPMDRMRSGFLISPQSMQPDITFAQDEEQKSQHQLQ  592

ScSte12       ------YPPSTFYSQN-INQYPRRRTVGMKSSQGNVPTGNKQSVGKSAKISKPLHIKTSAYQKQYKINLETKARPSAGDEDSAHPDKNKEISMPTPDSNTLVVQSEEGGAHSLEVDTNRR  679
NgSte12(1)    -------RQSQFDTRT-LGKRRKTKTCSKY----SSPNGIAR-------NTKPNKVLKNSINKMID-TLRSN---------------TKNVS----------------------------  598
NgSte12(2)    QHQHPNTHPPQLQHQLQHQRRPRTLRQGKYLKKS--------------------------I--------R--------------------------------------------------  628

ScSte12       SDKNLPDAT  688
NgSte12(1)    ---------  598
NgSte12(2)    ---------  628
```

**Supplementary Figure 2. Multiple sequence alignment of ScSte12, NgSte12(1), and NgSte12(2)**. The alignment was performed using Clustal Omega. A conserved phenylalanine is replaced by tyrosine (F-to-Y) in the DNA-binding motif of NgSte12(2). Additionally, the Dig1-binding motif is absent from both NgSte12(1) and NgSte12(2).

A

| PCR | Primer combination | Expected band (bp) |
|---|---|---|
| 1 | L641:TCCAGTCGCGTGACAATTTA<br>a_left520:ATTGAACCGCCAACTGATG | MATa: 1211 bp<br>MATalpha: none |
| 2 | L641:TCCAGTCGCGTGACAATTTA<br>a _left1622:AGGTAAATATGATAGACAGGGAGTGT | MATa: none<br>MATalpha: 981 bp |
| 3 | Yalpha 296:GCGAATTAGAGAAGAAAGGA<br>R425:GAATCCATCTGTGAGAATACC | MATa: none<br>MATalpha: 721 bp |
| 4 | Ya168:CTAACTTAGAACATACAGTTTGC<br>R425:GAATCCATCTGTGAGAATACC | MATa: none<br>MATalpha: 593 bp |

B

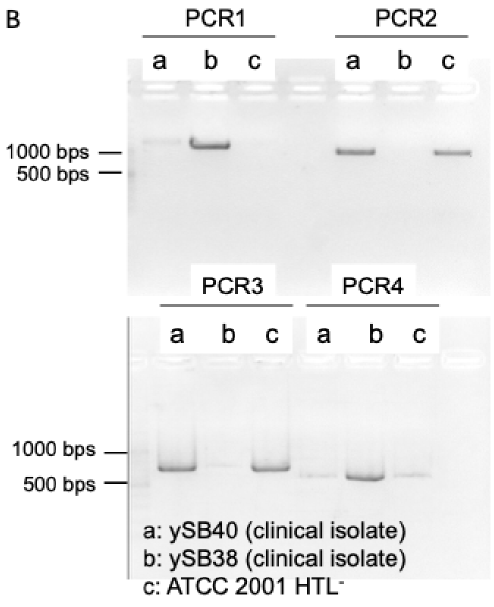


**Supplementary Figure 3. Mating type determination of *N. glabratus* ySB38 and ATCC2001 HTL⁻. A:** Primers used to determine mating type were extracted from [6] and the table lists primers for each PCR 1-4. **B.** PCR results for ySB38, ATCC2001 HTL⁻ and a second isolate ySB40.

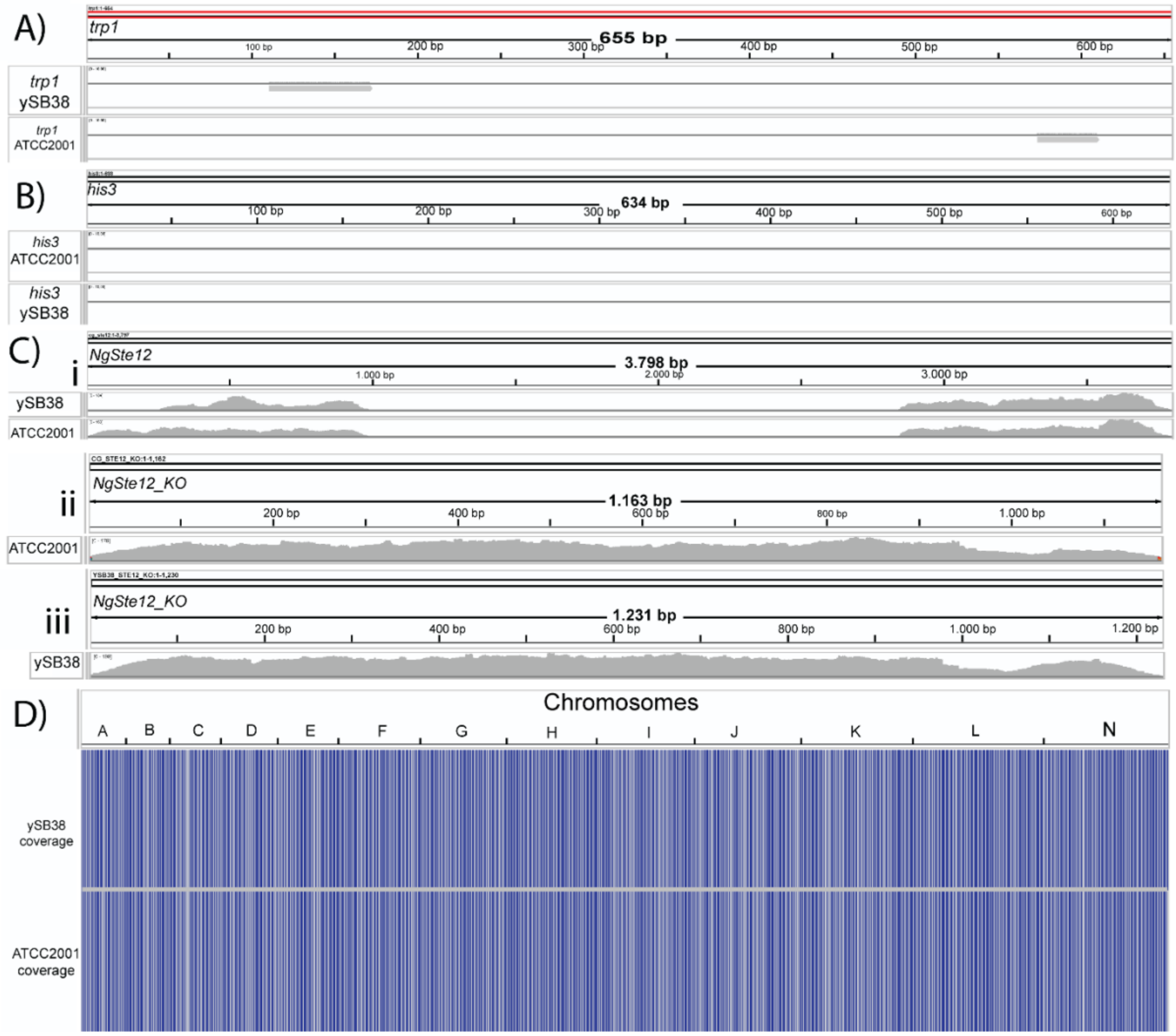


**Supplementary Figure 4. Aligned Illumina sequencing data indicate the absence of *trp1*, *his3*, and *ste12* in *N. glabratus* ATCC 2001ΔHTL⁻ and ySB98**. No reads align to the coding regions of **(A)** *trp1*, **(B)** *his3*, and **(C)** *ste12* (1) (i) despite high coverage of the adjacent regions. The created knockout strains contain the *nat* site in Ng *ATCC 2001HTL⁻* (ii) and a *nat-frt* site in ySB38 (iii) **(D)** Aligning sequencing data of both strains to the published reference genome of ATCC2001 results in high coverage of all regions. Blue heatmap regions indicate a minimum of 20x coverage.

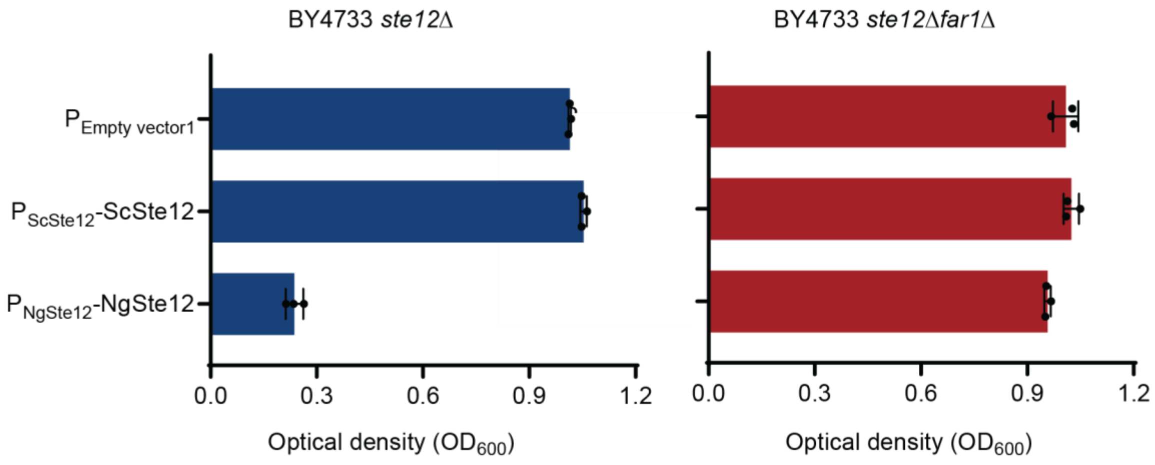


**Supplementary Figure 5. NgSte12(1) expression induces constitutive Far1-dependent growth arrest.** ScSte12 and NgSte12 under the control of their native promoters were cloned and transformed into BY4733 *ste12*Δ and *ste12*Δ*far1*Δ strains. Cells transformed with an empty vector were used as a control. All the strains were also transformed with a Venus reporter plasmid. Absorbance at 600 nm was measured after 20 hours in the absence of peptide. Data represent the mean ± SD of three biological replicates.

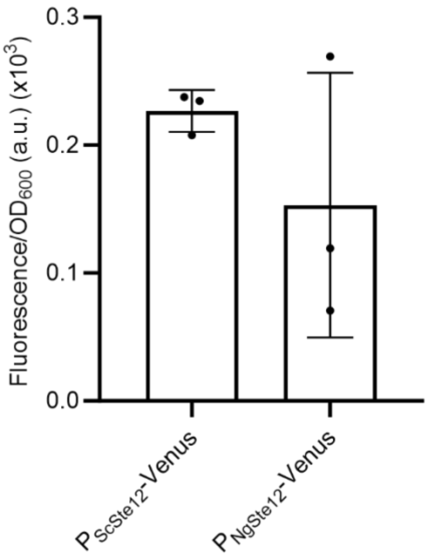


**Supplementary Figure 6. Fluorescence intensity driven by the ScSte12 and NgSte12 promoters** measured in *S. cerevisiae* BY4741 in the absence of peptide stimulation after 20 hours of growth. Autofluorescence of BY4741 transformed with an empty vector was provided as a reference baseline and subtracted from that induced by ScSte12 and NgSte12 promoter. Fluorescence was measured with an excitation wavelength of 488nm and an emission wavelength of 530nm and normalised to $OD_{600}$. Data represent the mean ± SD of three biological replicates.

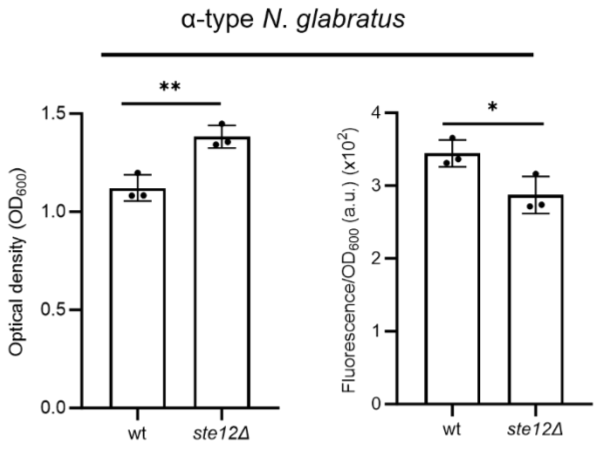


**Supplementary Figure 7. A Venus reporter under control of the NgFUS3 promoter was used to assess basal transcriptional activity of Ste12** in the absence of peptide-GPCR activation treatment in α-type *N. glabrata* mating types. Fluorescence was measured with an excitation wavelength of 488nm and an emission wavelength of 530nm and normalised to $OD_{600}$. Data represent the mean ± SD of three biological replicates.
